# Short- and long-term experience-dependent neuroplasticity interact during the perceptual learning of concurrent speech

**DOI:** 10.1101/2023.09.26.559640

**Authors:** Jessica MacLean, Jack Stirn, Alexandria Sisson, Gavin M. Bidelman

**Affiliations:** Department of Speech, Language, and Hearing Sciences, Indiana University, Bloomington, IN, USA; Program in Neuroscience, Indiana University, Bloomington, IN, USA; Cognitive Science Program, Indiana University, Bloomington, IN, USA

**Keywords:** Auditory perceptual learning, EEG, event-related brain potentials (ERP), frequency-following response (FFR), speech-in-noise perception

## Abstract

Plasticity from auditory experiences shapes brain encoding and perception of sound. However, whether such long-term plasticity alters the trajectory of short-term plasticity during speech processing has yet to be investigated. Here, we explored the neural mechanisms and interplay between short- and long-term neuroplasticity for rapid auditory perceptual learning of concurrent speech sounds in young, normal-hearing musicians and nonmusicians. Participants learned to identify double-vowel mixtures during ∼45 minute training sessions recorded simultaneously with high-density EEG. We analyzed frequency-following responses (FFRs) and event-related potentials (ERPs) to investigate neural correlates of learning at subcortical and cortical levels, respectively. While both groups showed rapid perceptual learning, musicians showed faster behavioral decisions than nonmusicians overall. Learning-related changes were not apparent in brainstem FFRs. However, plasticity was highly evident in cortex, where ERPs revealed unique hemispheric asymmetries between groups suggestive of different neural strategies (musicians: right hemisphere bias; nonmusicians: left hemisphere). Source reconstruction and the early (150-200 ms) time course of these effects localized learning-induced cortical plasticity to auditory-sensory brain areas. Our findings confirm domain-general benefits for musicianship but reveal successful speech sound learning is driven by a critical interplay between long- and short-term mechanisms of auditory plasticity that first emerge at a cortical level.

---

Experience on multiple timescales shapes our sensory systems. For example, studies have shown short-term changes in the selective tuning properties of cortical neurons that facilitates speech-in-noise (SIN) perception within a single training session (Ahveninen et al. 2011; Da Costa et al. 2013). Auditory perceptual learning studies also demonstrate rapid changes in behavior that track with subcortical (Carcagno and Plack 2011) and cortical (Ozaki et al. 2004) modulations as indexed by scalp-recorded event-related brain potentials (ERPs). Similarly, neuroimaging studies show long-term listening experiences (e.g., specific language expertise, bilingualism) shape the brain’s representations for speech sounds, selectively enhancing native compared to nonnative elements of a listener’s lexicon (Kuhl et al. 1992; Krishnan et al. 2010; Jeng et al. 2011; Krishnan et al. 2011; Bidelman and Lee 2015). Yet, because they are studied in isolation, the way in which short-term and long-term plasticity interact within the auditory system remains poorly understood.

Musicianship (e.g., formal music training) offers one approach to investigate brain plasticity along multiple timescales as many consider musicians to be a model of auditory neuroplasticity across the lifespan (Kraus and Chandrasekaran 2010; Herholz and Zatorre 2012; Alain et al. 2014; Moreno and Bidelman 2014). For example, trained musicians demonstrate improved behavioral performance in difficult listening scenarios, such as SIN and other “cocktail party” environments (Parbery-Clark, Skoe, Lam, et al. 2009; Coffey et al. 2017; Yoo and Bidelman 2019; Bidelman and Yoo 2020). On a smaller timescale, studies have demonstrated behavioral and neural encoding improvements in older adults’ speech and auditory processing following short-term music interventions (Alain et al. 2019; Dubinsky et al. 2019; Bidelman et al. 2022). Improvements to SIN perception following music training have also been observed in clinical populations, such as individuals with hearing loss (Lo et al. 2020). One explanation for musicians’ SIN benefit is the OPERA hypothesis, which posits that the precise auditory demands of music, when combined with repetition and high emotional reward, require attention-modulated engagement in overlapping networks for music and speech perception which confer benefits to speech processing (Patel 2011, 2014). Collectively, these studies suggest that long-term musical training (and related short-term interventions) bolster the brain’s ability to precisely encode speech sounds, which may confer advantages not only to everyday speech processing but also novel sound acquisition via learning (e.g., second language development; Slevc and Miyake 2006; Chobert and Besson 2013; Picciotti et al. 2018).

Musicians’ behavioral advantages for SIN processing are supported biologically through their stronger and faster sound-evoked neural responses at both brainstem and cortical levels of auditory processing (Shahin et al. 2003; Parbery-Clark, Skoe and Kraus 2009; Bidelman and Krishnan 2010; Bidelman et al. 2014). In particular, a large body of research has shown enhanced subcortical responses to sound with long-term music training, as measured via frequency-following responses (FFRs) (Musacchia et al. 2007; Carcagno and Plack 2011; Strait et al. 2012; Weiss and Bidelman 2015). At the cortical level, musicians show enhanced responses in the timeframe of the P2 wave of the ERPs, an early (150-200 ms) positive component reflecting perceptual sound object formation, speech identification, and concurrent speech segregation abilities (Shahin *et al*. 2003; Bidelman et al. 2013; Leung et al. 2013; Ross et al. 2013; Bidelman and Yellamsetty 2017). Presumably, such coordination in neural encoding across different levels of the auditory system might account for musicians’ superior ability to cope with real-world speech listening scenarios, including parsing target speech from background noise or competing talkers (Yoo and Bidelman 2019; Bidelman and Yoo 2020; Brown and Bidelman 2022).

One way to assess real-world SIN listening skills (and neuroplasticity therein) is via concurrent vowel identification tasks (Assmann and Summerfield 1989, 1990; Alain et al. 2007; Bidelman and Yellamsetty 2017). In these paradigms, listeners hear two simultaneous vowels and are asked to correctly identify both tokens as quickly and accurately as possible.

Behaviorally, speech identification accuracy improves with increasing pitch differences between vowels for fundamental frequency (F0) separations from 0 to about 4 semitones (Arehart et al. 1997; Chintanpalli and Heinz 2013; Chintanpalli et al. 2016). Critically, task success requires multiple processes: listeners must first segregate and then identify both elements of the speech mixture. The segregation of complex auditory mixtures is thought to reflect a complex, distributed neural network involving both subcortical and cortical brain regions (Palmer 1990; Sinex et al. 2002; Dyson and Alain 2004; Alain et al. 2005; Bidelman and Alain 2015). As such, double-vowel identification provides an ideal avenue for studying possible differential plasticity across the auditory system as it relates to learning and improving complex listening skills.

Using double vowel tasks, multiple studies have shown that short-term auditory perceptual learning (in nonmusicians) results in enhancements in the auditory cortical ERPs (Atienza and Cantero 2001; Reinke et al. 2003; Alain *et al*. 2007; Alain et al. 2015). The early timing of these neural changes (∼100-250 ms) suggests that learning induces reorganization of the sensory-receptive fields of auditory cortical neurons rather than procedural learning alone (Fritz et al. 2003; Alain *et al*. 2007). Similar short-term plasticity has been observed at a subcortical level in brainstem FFRs—though for isolated rather than double vowel speech sound training (Reetzke et al. 2018). However, training sessions in most of these studies took place over multiple days or even weeks, which probably involved longer-term mechanisms of learning (e.g., memory/sleep consolidation, overlearning) rather than sensory plasticity, *per se* (but see Alain *et al*. 2007). It is also unclear from the extant literature whether learning (i) engenders similar magnitudes of plasticity in auditory brainstem and cortex, and (ii) whether it proceeds in a bottom-up (e.g., brainstem→cortex) or top-down (e.g., cortex→brainstem) guided manner (cf. Ahissar and Hochstein 2004; Reetzke *et al*. 2018).

Here, we aimed to extend prior studies on the neural mechanisms of auditory perceptual learning by investigating the interplay between short- and long-term plasticity on concurrent speech processing. We build upon prior work by utilizing a concurrent double-vowel learning paradigm previously shown to induce rapid cortical plasticity in the ERPs (Alain *et al*. 2007). We measured simultaneous behavioral and multichannel EEG responses while listeners completed ∼45 minutes of training to assess short-term perceptual learning effects. The double-vowel paradigm provided an ideal test bed to assess changes in ecological speech listening skills. Our EEG approach also included the tandem recording of both brainstem FFRs with cortical ERPs to assess potential differential neuroplasticity at sub-vs. neocortical levels of the auditory-speech hierarchy. Comparisons between trained musician and nonmusician listeners allowed us to further assess the impact of long-term auditory experience on the trajectory of rapid perceptual learning. Our findings demonstrate brainstem and cortical levels of auditory processing are subject to different time courses of plasticity and reveal a critical interaction between long- and short-term neural mechanisms with regard to speech-sound learning.

## MATERIALS AND METHODS

### Participants

Twenty-seven young adults (ages 18-34; mean ± *SD*: 23.68 ± 4.22; 13 female) participated in this study. All participants had normal hearing thresholds bilaterally (pure tone average < 25 dB HL) at octave frequencies between 250 and 8000 Hz, were fluent in American English, and reported no history of neurologic or psychiatric disorders. Participants gave written, informed consent in accordance with a protocol approved by the Indiana University Institutional Review Board. Participants were recruited and separated into two groups based on their amount of music training. Musicians (M; *n* = 13) had ≥10 years of formal music training starting at or before age 12 (Wong et al. 2007). Nonmusicians (NM; *n* = 14) had ≤ 5 years of lifetime music training.

Groups did not differ on age (*t*(25) = 1.58; *p* = 0.413), cognitive ability as assessed through the Montreal Cognitive Assessment (Nasreddine et al. 2005) (*t*(25) =1.78; *p* = 0.088), self-reported bilingualism, (*X^2^*(1, *N* = 27) = 0.022, *p* = 0.883), sex balance (*X ^2^*(1, *N* = 27) = 1.78, *p* = 0.182), nor handedness (*t*(25) = −0.615; *p* = 0.544) (Oldfield 1971). As a confirmation of our group separation, musicians expectedly had ∼14 more years of music training than their nonmusician peers (M: 16.1 ± 4.3 years; NM: 2.4 ±1.7 years; *t*(25) = 10.93; *p* < 0.001).

### Double-vowel stimuli and task

Concurrent vowel stimuli were modeled after previous studies (Assmann and Summerfield 1989, 1990; Alain *et al*. 2007; Bidelman and Yellamsetty 2017). Stimuli consisted of synthesized, steady-state vowels (/a/, /e/, and /i/) which were presented in three unique vowel combinations (i.e., /a/ + /e/; /e/ + /i/; /a/ + /i/). Vowels were never paired with themselves. Stimuli were created with a Klatt-based synthesizer (Klatt 1980) coded in MATLAB (v 2021; The MathWorks, Inc., Natick, MA). Each vowel was 100 ms in duration with 10-ms cos^2^ onset/offset ramping to prevent spectral splatter. The fundamental frequency (F0) between vowels was set at 4 semitones (150 and 190 Hz), which promotes segregation for most listeners (Assmann and Summerfield 1990; Bidelman and Yellamsetty 2017). Importantly, the high F0s of both speech tokens was well above the phase-locking limit of cortical neurons and thus ensured FFRs were of a subcortical origin (Joris et al. 2004; Brugge et al. 2009; Bidelman 2018; Gorina-Careta et al. 2021). F0 and the first two formant frequencies (*F1*_a,e,i_ = 787, 583, 300 Hz; *F2* _a,e,i_ = 1307, 1753, 2805 Hz) remained constant for the duration of the token.

The speech sounds were presented in rarefaction phase through a TDT RZ6 interface (Tucker-Davis Technologies, Alachua, FL) controlled via MATLAB. Stimuli were presented binaurally at 79 dB SPL through electromagnetically shielded (Campbell et al. 2012; Price and Bidelman 2021) ER-2 insert earphones (Etymotic Research, Elk Grove, IL). Prior to EEG testing, we required all participants to identify single vowels with 100% accuracy. This ensured subsequent learning would be based on improvements in *concurrent* speech identification rather than isolated sound labeling, *per se*.

We used a clustered stimulus paradigm (Bidelman 2015) employing interspersed fast and slow interstimulus intervals (ISIs) to simultaneously collect brainstem FFRs and cortical ERPs during the active perceptual task. Each trial consisted of one of the three vowel combinations. During a trial, 20 repetitions of the vowel pair were presented with a fast ISI of 10 ms to elicit the FFR. The ISI was then slowed to 1100 ms and a single stimulus was presented to evoke the ERP. Participants then identified both vowels through keyboard responses following the isolated vowel pair. The next trial began after the participants’ response and 250 ms of silence. Participants were asked to identify both vowels as quickly and accurately as possible (no feedback was provided). Double vowel pairs were randomized in order. This identical task was repeated over four learning blocks. In total, each block generated 3000 FFR trials and 150 ERP trials. Each block took 10-15 minutes to complete. Participants were offered a short (2-3 min) break after each block to avoid fatigue.

### EEG recording and preprocessing

We used Curry 9 (Compumedics Neuroscan, Charlotte, NC) and BESA Research 7.1 (BESA, GmbH) to record and preprocess the continuous EEG data. Continuous EEGs were acquired from 64-channel Ag/AgCl electrodes positioned at 10-10 scalp locations (Oostenveld and Praamstra 2001). Recordings were digitized at 5 kHz using Neuroscan Synamps RT amplifiers. Data were referenced to an electrode placed 1 cm behind Cz during online recording. Data were re-referenced to linked mastoids (FFR) or common average reference (ERP) for subsequent analysis. Impedances were kept below 25 kΩ. Single electrodes were also placed on the outer canthi of the eyes and superior and inferior orbit to capture ocular movements. Eyeblinks were corrected using principal component analysis (Wallstrom et al. 2004). Responses were collapsed across vowel pairs to obtain an adequate number of trials for FFR/ERP analysis (Bidelman and Yellamsetty 2017; Yellamsetty and Bidelman 2018). Responses exceeding 150 µV were rejected as further artifacts. We then separately bandpass filtered the full-band responses from 120 to 1500 Hz and 1 to 30 Hz (zero-phase Butterworth filters; slope = 48 dB/octave) to isolate FFRs and ERPs, respectively (Musacchia et al. 2008; Bidelman *et al*. 2013; Price and Bidelman 2021). Data were then epoched (FFR: 0–105 ms; ERP: −200 – 1000 ms), pre-stimulus baselined, and ensemble averaged to obtain speech-evoked FFR and ERP potentials for each stimulus. To reduce the dimensionality of the data, we averaged waveforms for the three vowel pairs to collapse across tokens.

### FFR analysis

Brainstem responses were analyzed at the Cz electrode, where speech-FFRs are optimally recorded at the scalp (Bidelman 2015). From FFR waveforms, we computed the Fast Fourier Transform (FFT) for each block to analyze responses in the spectral domain. We measured the magnitude of the F0 response to both vowels, corresponding to the lower and higher vowel’s voice pitch (i.e., 140-160 Hz and 180-200 Hz). Prior literature has shown perceptual effects in speech-FFR are largely captured at the response F0 (Price and Bidelman 2021; Carter and Bidelman 2023; Rizzi and Bidelman 2023). The ±20 Hz search window was guided by the F0 of evoking double-vowel stimulus. Peak F0 amplitudes were averaged across vowels to derive a singular measure of FFR strength for each training block.

### ERP analysis

ERPs were analyzed at both the electrode and source level (Bidelman and Howell 2016). As with FFRs, scalp-level ERPs were quantified at the Cz electrode. To analyze the data at the source-level, we transformed each listener’s scalp potentials into source space using BESA’s Auditory Evoked Potential (AEP) virtual source montage (Scherg et al. 2002; Bidelman et al. 2018; Mankel et al. 2020). This applied a spatial filter to all electrodes that calculated their weighted contribution to the scalp recordings. We used a four-shell spherical volume conductor head model (Sarvas 1987; Berg and Scherg 1994) with relative conductivities (1/Ωm) of 0.33, 0.33, 0.0042, and 1 for the head, scalp, skull, and cerebrospinal fluid, respectively, and compartment sizes of 85 mm (radius), 6 mm (thickness), 7 mm (thickness), and 1 mm (thickness) (Picton et al. 1999; Herdman et al. 2002). The AEP model includes 11 regional dipoles distributed across the brain including bilateral auditory cortex (AC) [Talairach coordinates (*x,y,z*; in mm): *left* = (−37, −18, 17) and *right* = (37, −18, 17)]. Regional sources consist of three dipoles describing current flow (units nAm) in the three cardinal planes. We extracted the time courses of the tangential components for left and right AC sources as these orientations capture the majority of variance describing the auditory cortical ERPs (Picton *et al*. 1999). This approach allowed us to reduce each listener’s 64 channel ERP data to 2 source channels describing neuronal activity localized to the left and right hemisphere AC (Price et al. 2019; Mankel *et al*. 2020; Momtaz et al. 2021).

From sensor and source waveforms for each block, we measured the amplitude and latency of the P2 deflection between 130-170 ms. The analysis window was guided by visual inspection of the grand averaged data. We focus on the P2 as we have previously shown this neural index is sensitive to long-term plasticity of musicianship (Bidelman *et al*. 2014; Mankel *et al*. 2020), success in double-vowel identification (Bidelman and Yellamsetty 2017), and tracks with perceptual learning during auditory categorization tasks (Mankel et al. 2022; see also Alain et al., 2007).

### Statistical analysis

Unless otherwise noted, we analyzed the dependent variables using mixed-model ANOVAs in R (version 4.2.2) (R Core Team, 2020) and the *lme4* package (Bates et al. 2015). Fixed effects were block (4 levels; 1-4) and group (2 levels; musicians vs. nonmusicians). Subjects served as a random effect. Multiple comparisons were corrected via Tukey-Kramer adjustments. Effect sizes are reported as partial eta squared (η^2^_p_) and degrees of freedom (*d.f.*) using Satterthwaite’s method. Percent correct data were arcsine transformed to improve homogeneity of variance assumptions necessary for parametric ANOVA (Studebaker 1985). A priori significance level was set at α = 0.05.

We used repeated measure correlations (rmCorr, version 0.6.0) (Bakdash and Marusich 2017) to assess brain-behavior relationships within each group. Unlike conventional correlations, rmCorrs account for non-independence among observations (here, blocks within subjects) and measures within-subject correlations by evaluating the common intra-individual association between two measures.

## RESULTS

### Behavioral data

Figure 1 shows behavioral results across the four training blocks for both groups. RTs and identification accuracy were highly negatively correlated [*r* = −0.50, *p* < 0.001], indicating a typical time-accuracy tradeoff in concurrent vowel identification with improvements in learning (Bidelman and Yellamsetty 2017; Yellamsetty and Bidelman 2019). An ANOVA conducted on double-vowel identification accuracy (Fig. 1a) revealed a sole main effect of block [*F*(3, 75) = 12.13, *p* < 0.001, η^2^_p_ = 0.33]. The block effect was attributed to a steady increase in performance with training for both groups (linear contrast: M: *t*(75) = 4.34, *p* < 0.001; NM: *t*(75) = 3.01, *p* = 0.0035). In contrast, reaction times (Fig. 1b) were modulated by both block [*F*(3, 75) = 22.32, *p* < 0.001, η^2^_p_ = 0.47] and group [*F*(1, 25) = 4.49, *p* = 0.044, η^2^_p_ = 0.15]. The block effect was attributed to more rapid decision speeds at the end compared to beginning of training (M: *t*(75) = −4.76, *p* < 0.001; NM: *t*(75) = −5.43, *p* < 0.001). Yet, musicians showed faster RTs across the board. This was confirmed via correlational analyses, which showed listeners’ degree of musical training negatively predicted RTs; i.e., more highly trained individuals had faster decision speeds [*r* = −0.26, *p* = 0.008] (not shown). These data suggest that behaviorally, all participants improved in speed and accuracy with training, but musicians had overall faster response times to speech than nonmusicians.

**Figure 1.**
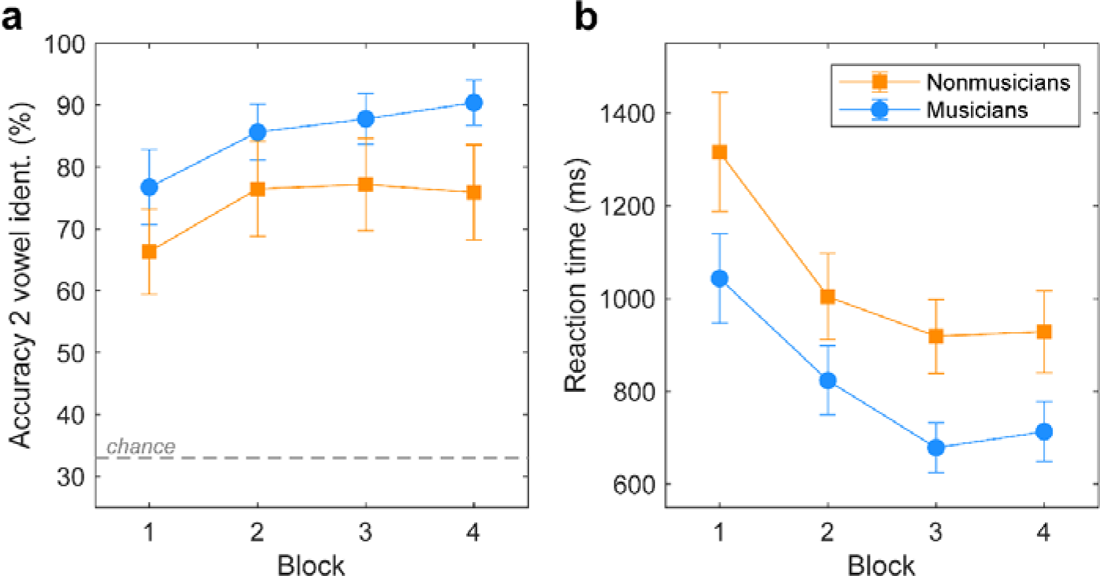
Behavioral performance across blocks show rapid perceptual learning of double-vowel speech stimuli. **(a)** Identification accuracy improved across blocks (each 10-15 min) for both groups. Chance performance on this task was 33%. **(b)** Reaction times improved across blocks for both groups but musicians were faster overall. Error bars = ± 1.s.e.m.

### Subcortical FFRs

Grand average FFR time waveforms and spectra are shown for each group and training block in Figure 2. FFRs showed phase-locked energy corresponding to the periodicities of both vowel stimuli. Response showed energy at the two F0s (i.e., 150 and 190 Hz) and their integer-related multiples up to the frequency ceiling of phase locking in the midbrain (∼1100 Hz) (Liu et al. 2006; Bidelman and Powers 2018). However, an ANOVA on FFR F0 amplitudes failed to reveal a block [*F*(3, 74) = 0.53, *p* = 0.66; η^2^_p_ = 0.02] or group effect [*F*(1, 25) = 0.42, *p* = 0.52; η^2^_p_ = 0.02]. These data suggest that brainstem speech representations were not modulated by either long- or short-term plasticity during rapid auditory perceptual learning.

**Figure 2.**
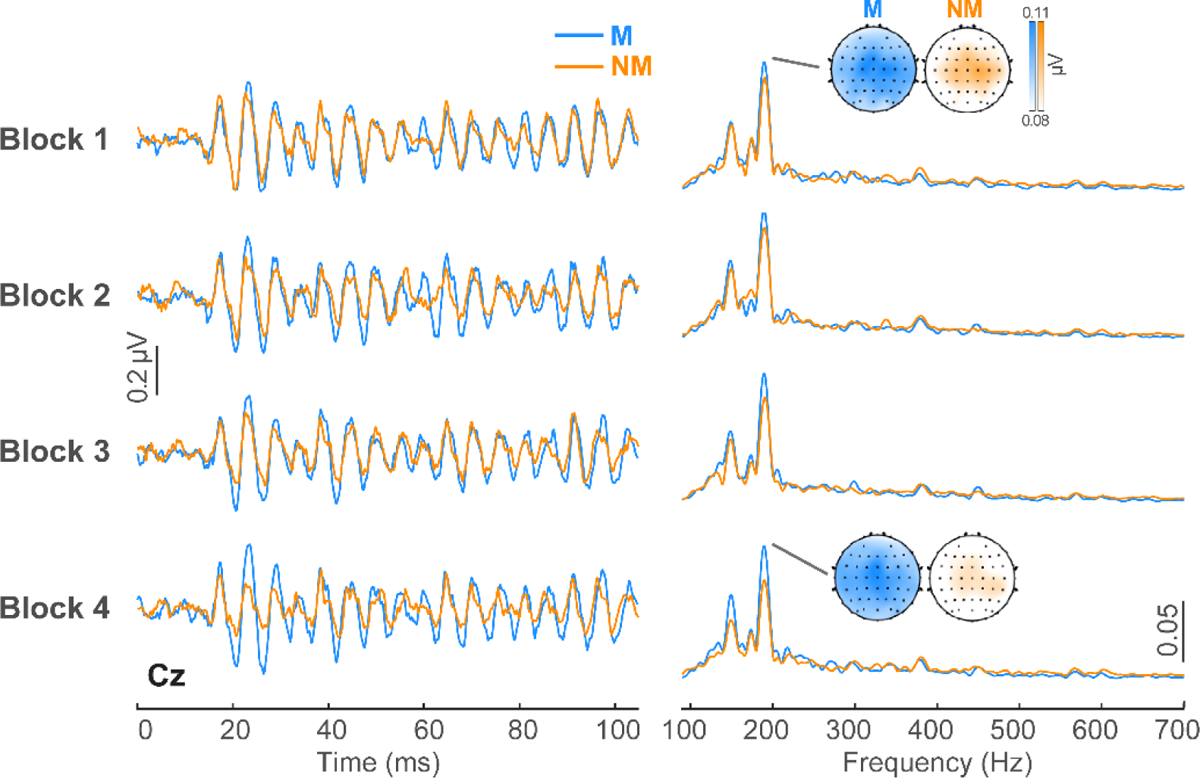
Subcortical FFR responses are not sensitive to rapid perceptual learning. FFR waveforms are shown across the four blocks for both groups in the time (left) and frequency (right) domains. Heatmaps show response strength at 190 Hz for Blocks 1 and 4, respectively. No group or block effects were observed in FFRs.

### Cortical ERPs

We analyzed cortical responses at the scalp and source levels. Figure 3a depicts scalp ERP responses at the electrode level (Cz) across the four blocks. Note prominent modulations in the P2 wave (∼150 ms) across training blocks. This deflection was maximal near the scalp vertex and inverted at the mastoids, consistent with generators in the supratemporal plane (Picton *et al*. 1999; Alain *et al*. 2007). No effects or interactions were observed for P2 latency (all *p*s > 0.07). However, an ANOVA conducted on P2 amplitudes revealed a block*group interaction [*F*(3, 74.2) = 3.88, *p* = 0.012; η^2^_p_ = 0.14) (Fig. 3b). Ms showed increased gains in P2 amplitude across blocks whereas NMs’ responses were invariant to learning (linear contrast: M: *t*(74) = 2.70, *p* = 0.009; NM: *t*(74) = −1.59, *p* = 0.12). Collectively, these data suggest a differential pattern in the strength of cortical speech encoding between groups (when measured at the scalp), with stronger training-related changes in neural processing among musically trained listeners. However, the volume-conducted nature of EEG does not allow us to adjudicate the intracranial generators underlying these effects in scalp data alone. Consequently, we used source analysis to determine if these neuroplastic changes observable in the sensor data were attributed to rapid changes *within* the auditory cortices themselves.

**Figure 3.**
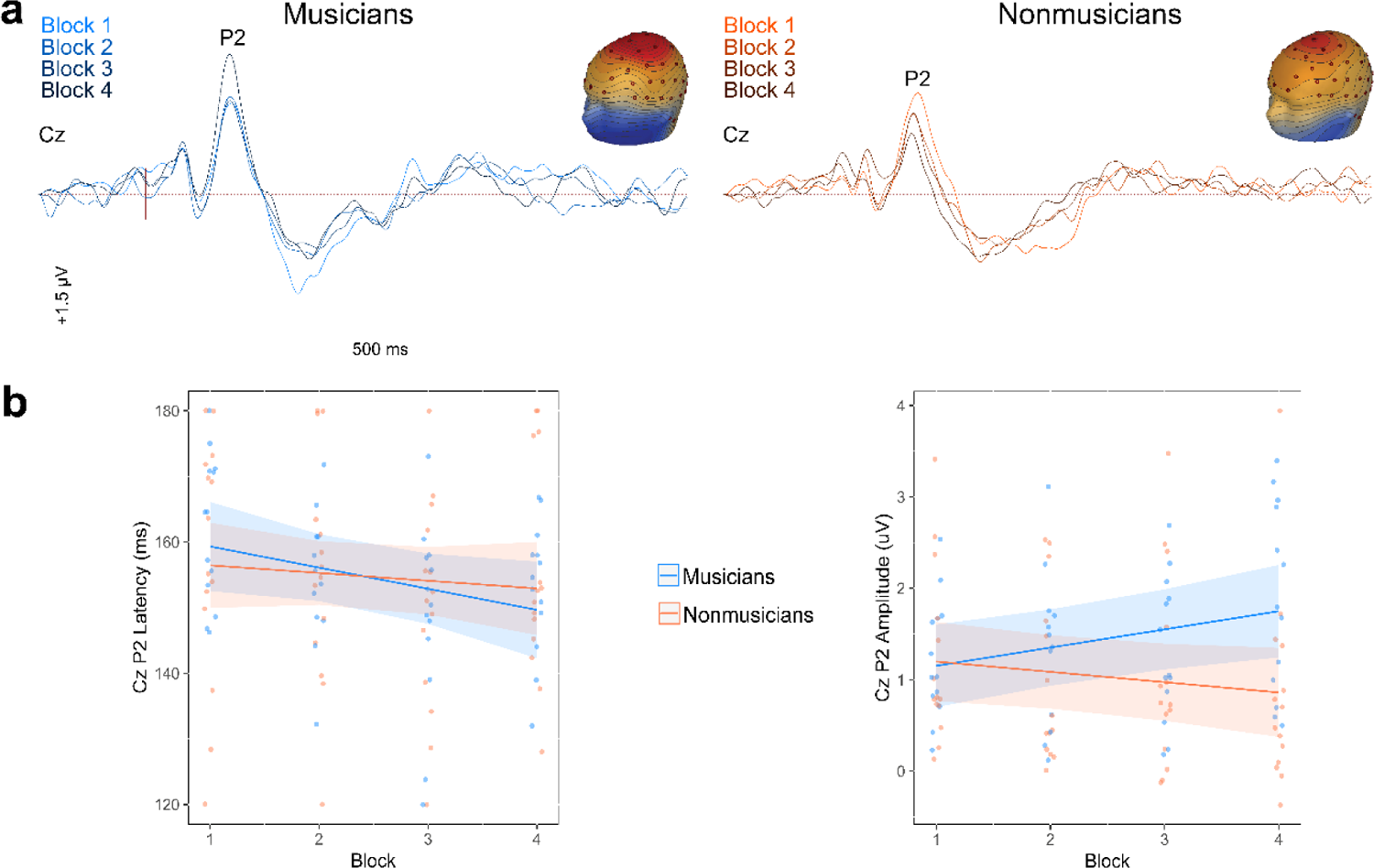
Cortical ERP responses (electrode level) are sensitive to rapid perceptual learning. **(a)** Grand average responses at Cz are displayed for both groups across the four blocks. Heatmaps show the scalp topography of the P2 (∼150 ms) for Block 4. **(b)** P2 amplitude and latency across blocks per group. Musicians showed increasing response strength with training whereas nonmusicians responses did not. Shading = ± 95% CI.

Figure 4 depicts grand average source responses for each group and a subset of blocks. Distributed imaging using Cortical Low resolution electromagnetic tomography Analysis Recursively Applied (CLARA; BESA v7.1) at the peak latency of the P2 (130-170 ms) (Iordanov et al. 2014) localized activity to bilateral AC (Fig. 4a). Source time courses from left and right hemisphere are shown in Figure 4b. We found training-related gains in P2 magnitude and latency that varied between groups and cerebral hemisphere (Fig. 5). While left hemisphere (LH) source magnitudes were invariant (all *p*s > 0.24; Fig. 5a), response latencies varied strongly with both training block [*F*(3, 73.5) = 5.28, *p* = 0.002; η^2^_p_ = 0.18] and group membership [*F*(1, 24) = 10.58, *p* = 0.003; η^2^_p_ =0.31]. The block effect was attributed to responses becoming progressively later with training (Fig. 5b) whereas the group effect was due to NMs having earlier (∼8 ms) responses compared to Ms overall.

**Figure 4.**
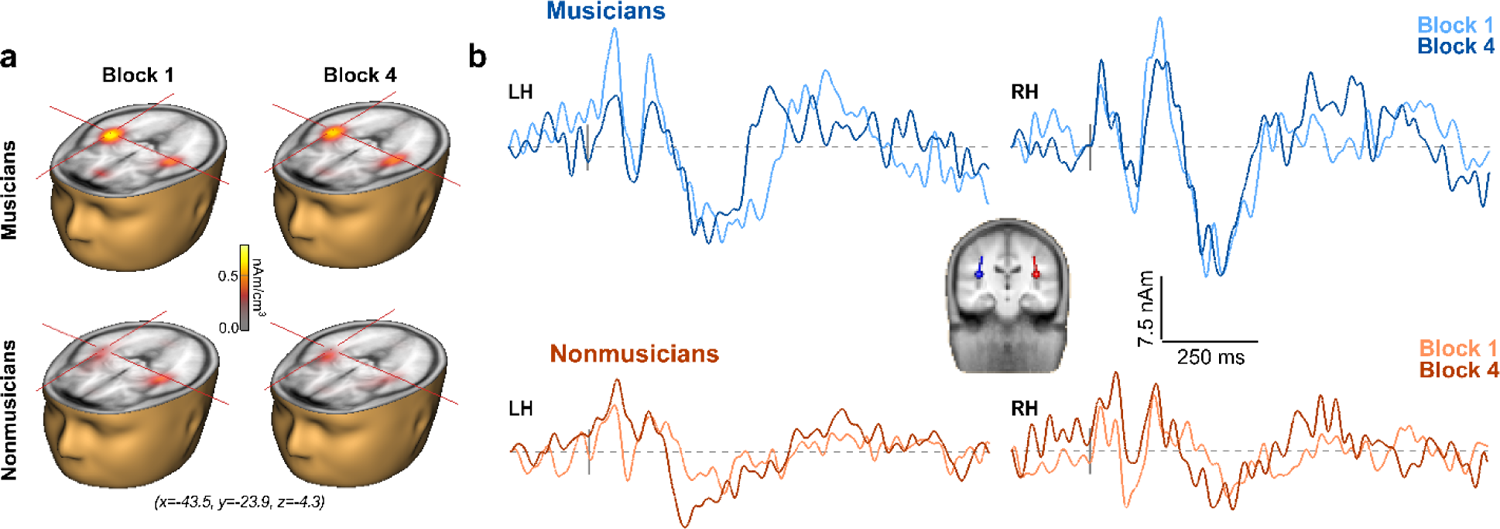
Speech coding in auditory cortex varies differentially with perceptual learning in musicians vs. nonmusicians. **(a)** Brain volumes show distributed source activation maps using CLARA imaging at the peak latency of the P2 (130-170 ms). Functional data are overlaid on the BESA brain template. The cross-hair demarks a representative voxel in right hemisphere primary auditory cortex (PAC; Talairach coordinates [cm]). **(b**) Source waveform time courses extracted from left (LH) and right (RH) hemisphere auditory cortex show different response trajectories as a result of learning between groups.

In contrast, right hemisphere (RH) P2 magnitudes showed a critical block*group interaction [*F*(3, 74.1) = 2.99, *p* = 0.036; η^2^_p_ = 0.11]. Whereas NMs’ RH response strength remained static across blocks [*t*(74) = 0.57, *p* = 0.57], Ms’ responses began more robust and became weaker with training to eventually converge with those of NMs [linear contrast: *t*(74.1) = −2.35, *p* = 0.022] (Fig. 5c). RH latencies revealed a sole main effect of group [*F*(1, 25.2) = 5.52, *p* = 0.027; η^2^_p_ = 0.18] that was again attributed to faster responses in NMs across the board (Fig. 5d). Collectively, these source data argue that the auditory cortex is sensitive to both perceptual learning for speech and prior listening experience (i.e., formal music training) which manifest in different lateralized effects in left vs. right hemisphere. In sum, we find that with speech-sound learning, auditory cortical processing (i) becomes prolonged in LH (though not made stronger/weaker) for all listeners; (ii) is slightly delayed in Ms overall; and (iii) is modulated in strength in RH among musicians.

**Figure 5.**
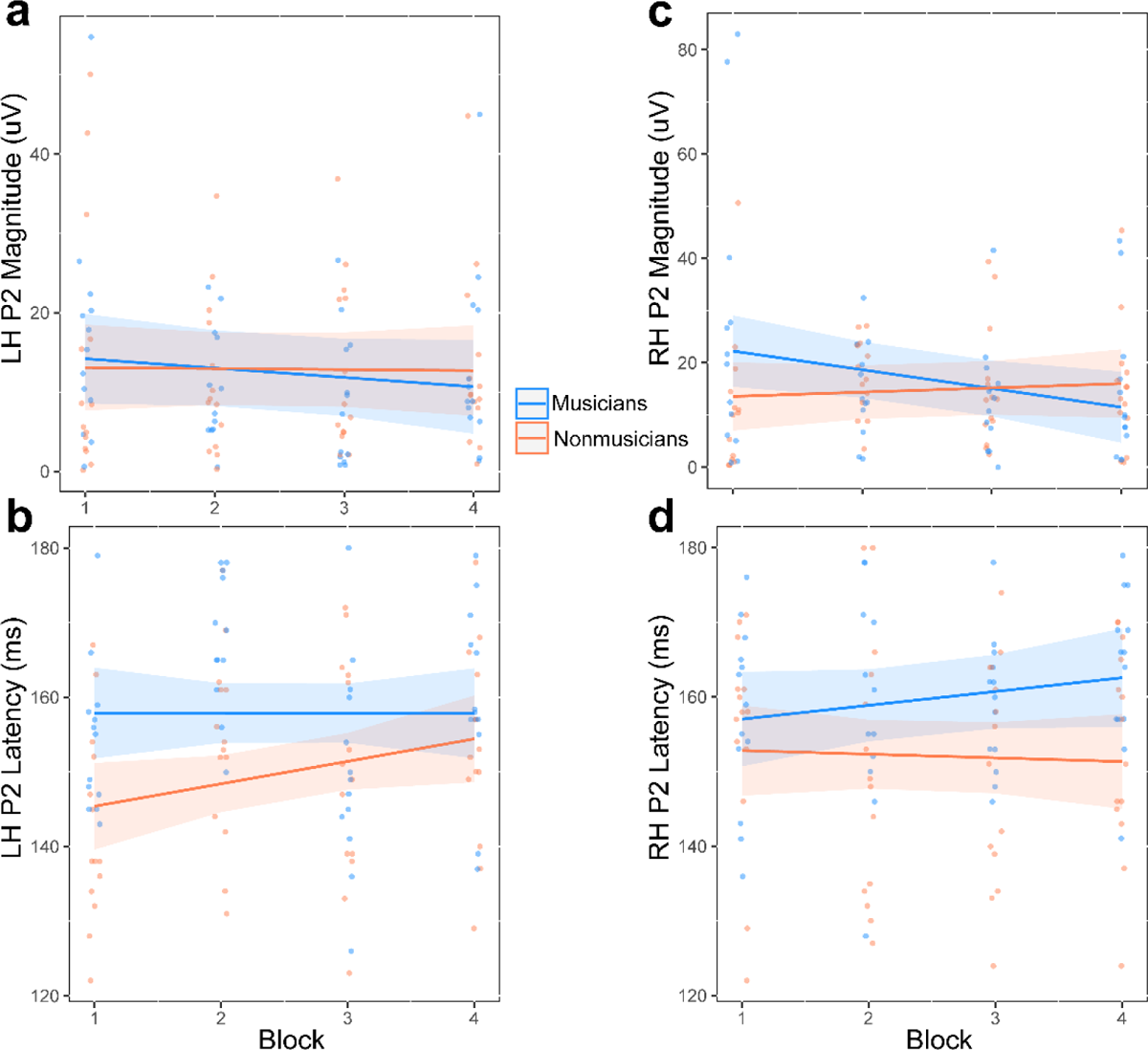
Latency and amplitude of source responses vary with group and hemisphere. **(a)** No effects were observed for LH P2 magnitude. **(b)** LH responses increased in latency with training, and NM had earlier responses than M overall. **(c)** RH magnitude remained static for NM, while M decreased across blocks. **(d)** NMs displayed earlier RH responses than Ms. Shading = ± 95% CI.

### Brain-behavior relationships

To assess correspondences between the neural and behavioral data, we performed repeated measure correlations (rmCorr) between RTs and source ERP amplitudes/latencies separately for Ms and NMs (Figure 6). We selected these measures given their sensitivity to group differences in our main analysis. We used rmCorr to account for the within-subject correlations stemming from the repeated testing across training blocks. Interestingly, we observed different trends between the two groups. In the left hemisphere, RT was negatively correlated with P2 latency for NMs [*r* = −0.38, *p* = 0.011], but not for Ms [*r* = −0.08, *p* = 0.61]. In the right hemisphere, RT was negatively correlated with P2 latency for Ms [*r* = −0.36, *p* = 0.025] but not for NMs [*r* = 0.19, *p* = 0.21]. No significant correlations were observed for RH P2 magnitude for either group [M: *r* = 0.26, *p* = 0.11; NM: *r* = −0.18, *p* = 0.26] (data not shown).

**Figure 6.**
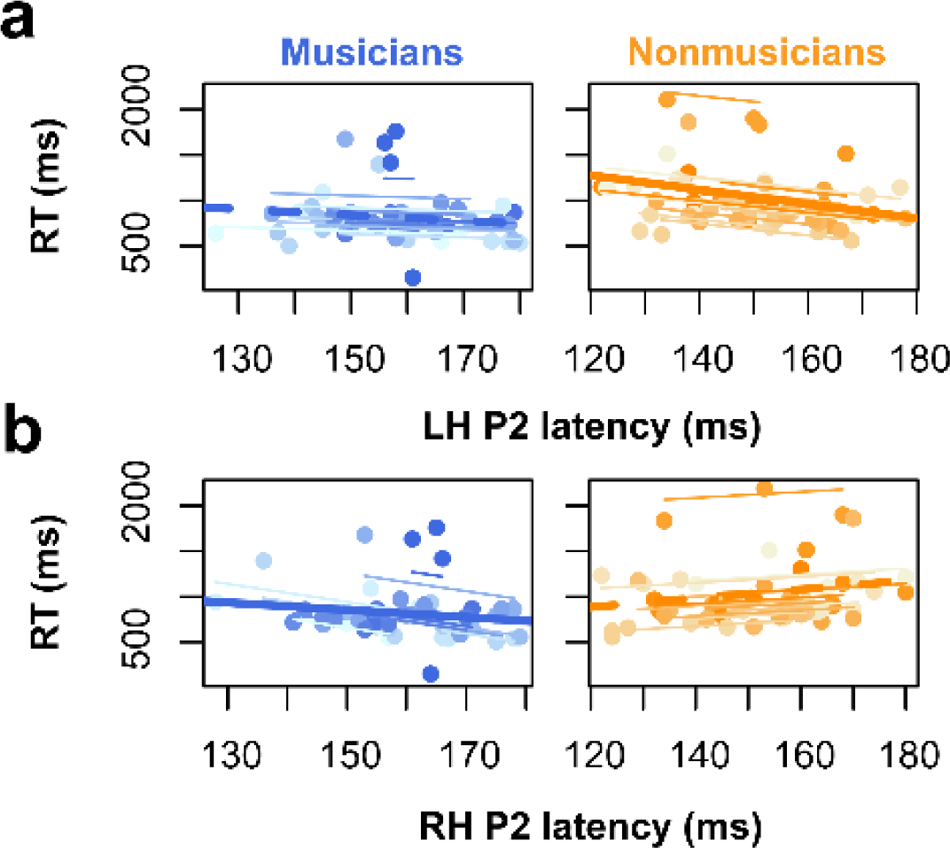
Correlations between neural and behavioral measures. Repeated measures correlations (rmCorr) (Bakdash and Marusich 2017) show the within-subject relation between measures for each listener (single points and thin lines) and overall association for the aggregate sample (thick lines). **(a)** RT was negatively correlated with LH P2 latency for NMs (but not for Ms). **(b)** RT was negatively correlated with RH P2 latency for Ms (but not for NMs). Solid trend lines = significant correlations; dotted trend lines = *n.s*.

## DISCUSSION

By simultaneously measuring behavior and EEG during a double-vowel learning task in musicians and nonmusicians, our data reveal three primary findings: (i) while both groups successfully learned to segregate speech mixtures, musicians were overall faster in concurrent speech identification than nonmusicians; (ii) short-term plasticity related to auditory perceptual learning for speech was not observed at the subcortical level; (iii) plasticity was highly evident at the cortical level where ERP responses revealed unique hemispheric asymmetries suggestive of different neural strategies between groups. Our findings demonstrate sub-vs. neocortical levels of auditory processing are subject to different time courses of plasticity and reveal a critical interaction between long- and short-term neural mechanisms with regard to speech-sound learning.

### Short-term plasticity from perceptual learning differs with long-term auditory experience

Our behavioral data replicate findings of Alain *et al*. (2007) demonstrating ∼45 min of training leads to rapid perceptual learning in deciphering speech mixtures. However, we extend these prior results by demonstrating faster overall reaction times for musicians relative to nonmusicians in double vowel identification across training blocks. This finding agrees with earlier data suggesting enhanced speech categorization in trained musicians (Elmer et al. 2012; Bidelman *et al*. 2014; Bidelman and Walker 2019) and highly musical novices (Mankel *et al*. 2020). Previous evidence suggests that musicianship automatizes categorical perception earlier in the auditory-linguistic brain networks subserving sound to label formation (Bidelman and Walker 2019), resulting in domain-general benefits applicable to speech processing. The faster behavioral responses observed here for musicians also aligns with previous findings (Schneider et al. 2002; Chartrand and Belin 2006; Bidelman *et al*. 2014; Bidelman and Walker 2019) by suggesting that stronger category representation in early auditory cortical structures might dictate the speed and trajectory of how listeners acquire speech labels during novel learning. This notion is further bolstered by the correlations we find between listeners’ RTs and source responses localized to primary auditory cortex.

Several previous studies have shown benefits of musical training for “cocktail party” speech listening tasks (Bidelman and Krishnan 2010; Parbery-Clark et al. 2011; Swaminathan et al. 2015; Clayton et al. 2016; Coffey *et al*. 2017; Deroche et al. 2017; Du and Zatorre 2017; Mankel and Bidelman 2018; Torppa et al. 2018; Yoo and Bidelman 2019; Maillard et al. 2023). The double vowel identification task used here serves as an extension of cocktail-party listening as it requires accurate identification of concurrent speech tokens. We found superior RTs (but not accuracy) in concurrent speech identification among musicians. The fact that group benefits were limited to speed may be due to the relative simplicity of our task. While accuracy was not at ceiling, participants’ performance was largely successful (>70%) even in the first training block. Concomitant neural changes notwithstanding, it is possible our task may not have “taxed” the perceptual system enough to elucidate strong behavioral differences beyond those seen in RT speeds. Nevertheless, our perceptual data alone demonstrate a domain-general benefit of music experience on the sound-to-label mapping process inherent to speech perception (Patel and Iversen 2007).

### Neural correlates of auditory perceptual learning are absent in subcortex

In stark contrast to the cortical ERPs (present study; Alain *et al*. 2007), 45 min of training failed to induce rapid plasticity in brainstem FFRs. This suggests sensory enhancements in brainstem auditory processing are neither necessary nor sufficient to yield learning-related reorganization in auditory cortex over the same (short) time course of training (present study; Alain *et al*. 2007). This notion aligns with recent studies suggesting cortical changes precede those in brainstem (Reetzke *et al*. 2018; Skoe et al. 2021) by several days/weeks, as well as theoretical accounts that learning proceeds in a top-down guided manner (Ahissar and Hochstein 2004), with sensory change in the brainstem only emerging at expert stages of learning (Reetzke *et al*. 2018).

Our data also failed to reveal FFR differences between musicians and non-musicians. This contrasts several studies that have shown enhanced F0-pitch encoding in speech-FFRs of musicians (Musacchia *et al*. 2007; Bidelman, Gandour, et al. 2011; Bidelman, Krishnan, et al. 2011; Bidelman *et al*. 2014; Coffey et al. 2016), though not always consistently (e.g., see Strait *et al*. 2012; Bidelman and Alain 2015). Indeed, at the behavioral level, musicians are not always better at exploiting F0 cues for voice segregation than their nonmusician peers (Deroche *et al*. 2017). The surprising lack of FFR group differences in the present data could be due to the relatively high F0 of our stimuli (> 150 Hz). High F0s minimize cortical contributions to the FFR (Bidelman 2018) and result in a dominantly brainstem-centric response that largely reflects exogenous processing of double-vowel stimuli (Yellamsetty and Bidelman 2019). Previous studies that have found musician encoding advantages and learning-related effects in the FFR have also used much lower F0s (∼100 Hz) (e.g., Song et al. 2008; Carcagno and Plack 2011; Chandrasekaran et al. 2012; Reetzke *et al*. 2018) which may have reflected cortical rather than subcortical plasticity, *per se*. It is also possible that auditory plasticity is stronger and emerges earlier at cortical relative to brainstem levels (e.g., Reetzke *et al*. 2018; Skoe *et al*. 2021; Bidelman *et al*. 2022; Lai et al. 2022) and varies in a stimulus-specific manner (Holmes et al. 2018). Nascent changes in the brainstem FFR might therefore require more protracted training than the short learning tasks used here. Indeed, lasting changes in the neural differentiation of speech, as indexed by the FFR, are observable no earlier than several days of training (Song *et al*. 2008; Reetzke *et al*. 2018) and even 1 year by some accounts (Kraus et al. 2014).

Additionally, prior studies demonstrating musician F0 benefits have exclusively used passive listening tasks. Attention varies with musical training (Strait et al. 2010; Yoo and Bidelman 2019) and is known to enhance the speech FFR (Price and Bidelman 2021; Lai *et al*. 2022; Carter and Bidelman 2023). Consequently, experience-dependent effects of music on FFR strength might be more muted under states of active attentional engagement if nonmusicians deploy more attentional resources during speech processing. This notion is indeed supported by the longer RTs and more invariant cortical P2 across blocks we find in nonmusicians, which suggest our behavioral task was more demanding and/or recruited more attentional resources in this group. Even so, participants were able to accomplish our task early in the training regimen, and therefore change in sensory representation at an early subcortical level was perhaps not necessary for task success. Instead, we observe salient changes in RTs and ERP neural timing which suggest that learning in our task probably reflected improved *access to,* rather than strength of, sensory representation at a cortical level of processing (Binder et al. 2004; Bidelman *et al*. 2014).

### Neural correlates of perceptual learning are robust in cortex and reflect different neural strategies between musicians and nonmusicians

Both our sensor and source ERP data were consistent in showing stronger learning-related neural changes in musicians. In particular, hemispheric PAC activity suggested distinct neural strategies with learning based on prior auditory experience: behavioral RTs correlated with right hemisphere P2 latency for musicians but left hemisphere P2 latency for nonmusicians. Given musicians’ faster overall behavioral speed in our task, the double-dissociation in hemispheric latencies between groups may point to superior “cue-weighting” by musicians towards pitch-related cues (Zatorre et al. 1992). Relationships between behavioral decision speeds and right hemispheric learning patterns for musicians could reflect their focus on pitch interval features (i.e., “musical” content) between the two vowels, whereas nonmusicians’ left hemispheric pattern may reflect heavier reliance on linguistic information (Mankel *et al*. 2022). Though our task relies on segregation and identification of speech tokens, musicians’ stronger learning-related changes in right hemisphere may indicate a focus on pitch rather than linguistic information of the speech stimuli (Alain *et al*. 2005), indicating distinct task strategies as a result of long-term experience in music.

A rightward biased mechanism in musicians is additionally supported by their decreased RH P2 magnitude with training. Declines in P2 strength with learning is consistent with other single-session, short-term learning experiments in which sensory-evoked neural responses become more efficient during active task engagement (Guenther et al. 2004; Alain et al. 2010; Ben-David et al. 2011; Pérez-Gay Juárez et al. 2019; Mankel *et al*. 2022). Stronger engagement of RH may also indicate increased attention to frequency-related information (Crowley and Colrain 2004).

P2 latency, especially in RH, better differentiates phonetic speech categories in nonmusicians with higher musicality (Mankel *et al*. 2020), paralleling our findings here in trained musicians. Our results are also consistent with prior neuroimaging work suggesting musicians and nonmusicians process speech categories by differentially engaging different nodes of the auditory-linguistic network for otherwise identical perceptual tasks. For example, whereas musicians regulate speech coding to relatively early auditory areas (e.g., PAC), nonmusicians recruit additional downstream brain mechanisms (e.g., inferior frontal regions; Broca’s area) to decode the same speech category labels (Bidelman and Walker 2019).

Somewhat surprisingly, we found learning-related changes in P2 timing were *negatively* correlated with behavioral RTs (in both groups though in opposite cerebral hemispheres). That is, later neural responses in PAC predicted faster speeds in double-vowel identification. The direction of this effect is not immediately apparent, as we would have expected faster RTs to correlate with earlier P2 responses. With respect to the direction of P2 modulation with learning, the literature has been somewhat equivocal. Different experiments have reported changes in evoked response in seemingly opposite directions (Tremblay et al. 2001; Atienza et al. 2002; Bosnyak et al. 2004; Sheehan et al. 2005; Zhang et al. 2005; Tong et al. 2009; Alain *et al*. 2010; Ben-David *et al*. 2011; Carcagno and Plack 2011; Ross *et al*. 2013; Wisniewski et al. 2020). It is possible that the counterintuitive earlier cortical responses—as we observe in nonmusicians— reflect increased arousal during task engagement. Indeed, RTs follow a U-shape with changes in arousal level such that they are fastest at intermediate levels and deteriorate (slow) in overly relaxed or tense states (Broadbent 1971; Welford 1980). Similarly, earlier P2 latency has been associated with more aroused and wakeful states of attention (Crowley and Colrain 2004).

Consequently, it is possible NMs were more taxed during the rapid speech identification task leading to increased arousal that manifested in their longer behavioral RTs and counterintuitively earlier P2 responses. Variations in arousal might also explain the negative RT-P2 relations we find in both groups. However, we also note the P2 itself reflects multiple sources with subcomponents in Heschl’s gyrus, planum temporale, and surrounding auditory associations in both hemispheres (Steinmetzger and Rupp 2023). Thus, it is also possible the hemispheric differences we find in RT-P2 relations between groups reflect unique engagement of these multiple P2 generators that are not captured by our single dipole foci.

### Interplay between short- and long-term plasticity in early auditory cortex

The P2 component of the ERPs occurs relatively early in the auditory cortical hierarchy (∼150 ms after stimulus onset). The learning-related changes seen in our data agree with previous studies showing associations between P2 and speech discrimination (Alain *et al*. 2010; Ben-David *et al*. 2011), sound object identification (Leung *et al*. 2013; Ross *et al*. 2013), and early speech category representation (Bidelman *et al*. 2013; Bidelman and Lee 2015; Alho et al. 2016; Bidelman and Walker 2019; Mankel *et al*. 2020). Interestingly, we show differential trajectories of neuroplasticity in sound encoding with rapid learning as a function of previous auditory experience. Musicians responded faster to vowel pairs and displayed greater learning-related changes compared to nonmusicians’ at a cortical level. This suggests that the long-term auditory experience of musicianship might act as a catalyst for novel sound learning that would be highly relevant in other domains (e.g., second language learning; Slevc and Miyake 2006; Chobert and Besson 2013; Picciotti *et al*. 2018). We argue the early nature of these effects in waves that localize to auditory-perceptual areas (and well before motor responses) suggests the rapid plasticity we observe with auditory learning can be attributed to changes in sensory encoding rather than later procedural learning (Alain *et al*. 2007; Mankel *et al*. 2022). Indeed, neither task familiarity (i.e., procedural learning) nor stimulus repetition alone are sufficient to produce changes in the early cortical ERPs (Alain *et al*. 2007; Mankel *et al*. 2022).

Despite clear musician advantages at behavioral and cortical levels, our speech learning task was not sufficient to observe subcortical changes which have been observed in longer training regimens over several sessions and days (Song *et al*. 2008; Carcagno and Plack 2011; Reetzke *et al*. 2018). Future studies using more difficult tasks and longer learning paradigms should be conducted to determine the dosage of training needed to induce learning-related plasticity at brainstem vs. cortical levels of the auditory pathway (cf. Reetzke *et al*. 2018), along with the effects of consolidation to learning gains (cf. Alain *et al*. 2015). More broadly, understanding the differential timelines in plasticity for speech coding resulting from musicianship would further support the use of music-based interventions to enhance speech and language outcomes.

## Acknowledgements

The authors thank Rose Rizzi for assistance in data collection and comments on earlier version of the manuscript. Requests for data and materials should be directed to G.M.B. This work was supported by the National Institute on Deafness and Other Communication Disorders (R01DC016267 to G.M.B.).

